# Changes in protonation states of in-pathway residues can alter ligand binding pathways obtained from spontaneous binding molecular dynamics simulations

**DOI:** 10.1101/2022.04.30.490145

**Authors:** Helena Girame, Marc Garcia-Borràs, Ferran Feixas

**Author notes:** Correponding Authors.

## Abstract

Protein-ligand binding processes often involve changes in protonation states that can be key to recognize and orient the ligand in the binding site. The pathways through which (bio)molecules interplay to attain productively bound complexes are intricate and involve a series of interconnected intermediate and transition states. Molecular dynamics (MD) simulations and enhanced sampling techniques are commonly used to characterize the spontaneous binding of a ligand to its receptor. However, the effect of protonation state changes of in-pathway residues in spontaneous binding MD simulations remained mostly unexplored. Here, we used molecular dynamics simulations to reconstruct the trypsin-benzamidine binding pathway considering different protonation states of His57. This residue is part of the trypsin catalytic triad and is located more than 10 Å away from Asp189, which is responsible for benzamidine binding in the trypsin S1 pocket. Our MD simulations showed that the binding pathways that benzamidine follow to target the S1 binding site are critically dependent on the His57 protonation state. Binding of benzamidine frequently occurs when His57 is protonated in the delta nitrogen while the binding process is significantly less frequent when His57 is positively charged. Constant-pH MD simulations retrieved the equilibrium populations of His57 protonation states at trypsin active pH offering a clearer picture of benzamidine recognition and binding. These results indicate that properly accounting for protonation states of distal residues can be important in spontaneous binding MD simulations.

## INTRODUCTION

Characterizing the mechanisms of ligand binding and unbinding to a biomolecule is crucial to elucidate the molecular basis of biological processes and improve the potency of drugs.^1^ The pathways through which (bio)molecules interplay to attain stable (and often transient) bound complexes are intricate and involve a series of interconnected intermediate, misbound, and transition states. Molecular dynamics (MD) simulations and enhanced sampling techniques are frequently used to characterize the spontaneous binding pathways of drugs, substrates, or peptides to its biological receptors.^2–4^ In these simulations, one or more ligands are commonly placed in the solvent and are allowed to freely diffuse without biasing the MD simulation toward a particular protein region. Providing sufficient simulation time and an accurate description of the system, the ligand freely explores the dynamic protein surface until it spontaneously finds its presumed binding site.^5^ Using long-time scale MD simulations, it was possible to reconstruct step by step the binding pathway of agonist (+)-pentazocine into the σ_1_ receptor.^6^ Markov-State Models were used to completely reconstruct ligand binding and unbinding pathways and the associated kinetics of enzyme-inhibitor complexes.^7,8^ Unconstrained enhanced sampling methods were used to simulate binding of allosteric modulators into G-protein coupled receptors^9^ or substrate binding in allosterically regulated enzymes.^10^ The predictive power of spontaneous binding simulations relies on being able to access the timescale required to sample the binding event and critically depends on the accurate description of the simulated system.

Around 60% of protein-ligand binding events involve changes in protonation states.^11,12^ Properly accounting for protonation states of protein residues is crucial to characterize ligand binding with MD simulations. The prediction of protonation states from rigid X-ray structures can lead to their incorrect assignment as even subtle structural fluctuations can affect each residue environment. With constant-pH molecular dynamics (CpH-MD) it is possible to model pH effects retrieving the protonation equilibriums of titratable residues coupled to protein conformational dynamics.^13–16^ Recently, Vo and co-workers reconstructed how fentanyl binds µ-opiod receptor with CpH-MD showing that the protonation of His257 at the binding pocket plays a crucial role to properly orient fentanyl.^17^ These results point out the importance of correctly accounting for protonation states of residues in the binding pocket to characterize the thermodynamics and kinetics of ligand-binding. However, as captured by spontaneous binding MD simulations, the ligand can establish contact with different protein residues in its pathway toward the binding site. The nature of these interactions will also be determinant for the kinetics of the ligand binding process. Despite the number of studies of protein-ligand pathways, the effect of protonation state changes of in-pathway residues in spontaneous binding simulations remains mostly unexplored.

The binding of benzamidine to trypsin has been commonly used as an enzyme-inhibitor model system for studying spontaneous binding and benchmarking enhanced sampling techniques due to the rapid formation of the trypsin-benzamidine complex.^5^ Trypsin is a serine protease responsible of hydrolyzing proteins through a catalytic triad formed by Ser195, His57, and Asp102 residues (see Figure 1). The positively charged inhibitor benzamidine is recognized in the specific S1 pocket which contains a negatively charged Asp189 located more than 10 Å away from the catalytic triad. In a landmark publication, Buch et al. reconstructed the free-energy landscape of benzamidine binding from a total of 495 MD simulations of 100 ns, observing productive binding in 38% of the simulations.^7^ By analyzing the independent trajectories, they observed that catalytic His57 and Ser195 residues were commonly found in the binding pathway of benzamidine in its way toward the S1 pocket. Plattner and Noé extended the previous MD simulations gathering a total of 150 μs of simulation time and identified seven interconnected metastable states of trypsin that present additional competitive binding sites that could not be revealed by means of experimental techniques.^8^ The binding of benzamidine have also been studied using unconstrained enhanced sampling methods by Miao and co-workers who reconstructed binding and unbinding pathways using Gaussian accelerated molecular dynamics (GaMD) simulations.^18^ Interestingly, unbinding pathways showed that benzamidine passes next to His57 in its dissociation from the S1 pocket to the solvent. Therefore, His57 play a prominent role in both catalysis and binding.

**Figure 1.**
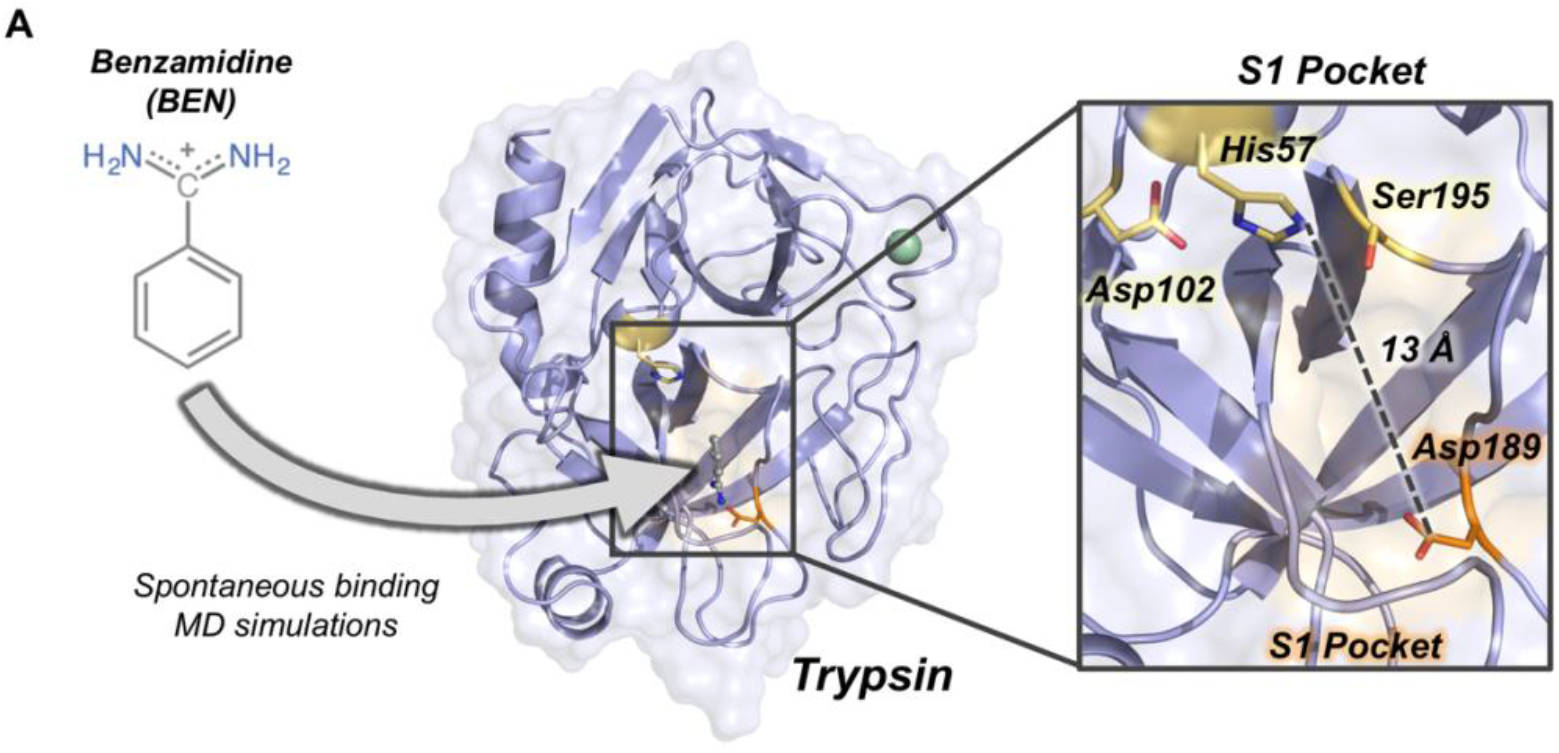
Overview of trypsin structure and S1 pocket. (A) Trypsin (in purple) structure corresponding to PDB ID 3PTB with benzamidine inhibitor (in grey) bound to the S1 pocket (orange surface). His57 and Asp189 residues are highlighted in yellow and in orange sticks, respectively. Calcium ion is depicted as a green sphere. Overview of the S1 pocket with catalytic residues (Asp102, His57, and Ser195) shown in yellow sticks. The distance between the carbon of the side-chain of Asp189 and the side-chain of His57 is 13 Å.

Enzymes are sensitive to pH changes and catalytic residues commonly change their protonation states at different stages of the catalytic cycle. Trypsin is active in a pH range between 7.0 and 9.0 and His57 is postulated to alter between two protonation states along the reaction mechanism.^19^ Czodrowski and co-workers studied the protonation changes in ligand binding in trypsin concluding that His57 is responsible for the most relevant pKa shifts during binding and catalysis.^20^ Most common software to assign protonation states from X-ray structures predict a positively charged His57 (both delta and epsilon nitrogens protonated, HIP) at pH = 7.0 while a less clear picture arises at pH = 8.0, where both HIP and neutral His57 with the delta nitrogen protonated (HID) are possible protonation states. Short-time scale MD simulations revealed that both HIP and HID were possible protonation states of His57.^21^ Spontaneous binding simulations of the benzamidine-trypsin system have been commonly performed with His57 in the HID state, which is the assumed protonation state when the Michaelis complex is formed.^22^ The question is whether the protonation state of His57 can influence benzamidine binding.

Here, we use spontaneous binding MD simulations to reconstruct the trypsin-benzamidine binding pathway considering different protonation states of His57. This histidine is part of the trypsin catalytic triad and is located more than 10 Å away from Asp189 responsible for benzamidine binding in the S1 pocket (Figure 1). Our MD simulations show that the spontaneous binding pathways are critically dependent on the His57 protonation state. Binding of benzamidine frequently occurs in a few hundreds of nanoseconds when histidine is protonated in delta (HID) while productive binding is scarcely observed when His57 it is positively charged (HIP). CpH-MD simulations reflect that both HID and HIP forms are significantly populated at the pH range between 7.0 and 8.0 showing the displacement of the equilibrium toward the HID protonation state upon pH increase. These results indicate that properly accounting for protonation states of distal residues can be key to obtain reliable pathways in spontaneous binding simulations.

## METHODS

### System Preparation

We used the crystal structure of benzamidine-bound *Bos taurus* trypsin (PDB ID 3PTB) as starting point for our molecular dynamics (MD) simulations. First, benzamidine was removed from the S1 pocket to protonate the system. Second, protonation states of all protein residues were assigned based on 3.0 H++ webserver (http://biophysics.cs.vt.edu/H++) at pH 7.0.^23^ H++ suggests that His57 is positively charged (HIP) at both pH = 7.0 and 8.0. On the other hand, PROPKA 3^24^ assigns His57 as HIP at pH = 7.0 and as neutral, with the delta nitrogen protonated (HID), at pH = 8.0. To explore the role of His57 protonation in benzamidine spontaneous binding, we manually assigned the protonation of His57 residue to either HID, HIE, or HIP. Once protonated, four benzamidine molecules were arbitrarily placed in the solvent, 30 Å away from Asp189 binding pocket. Parameters for MD simulations for benzamidine were obtained from the generalized AMBER force field (GAFF),^25^ with partial charges set to fit the electrostatic potential generated at HF/6-31G* level of theory by restrained electrostatic potential (RESP) model.^26^ The atomic charges were calculated according to the Merz−Singh−Kollman^27,28^ scheme using Gaussian 16 (M. J. Frisch et al, 2016).

### Conventional Molecular Dynamics Simulations

Spontaneous MD simulations starting from three different protonation states of His57 were performed in explicit water using AMBER18 package.^30^ AMBER-ff14SB force field^31^ was used to describe the protein, GAFF^25^ for benzamidine, and TIP3P for water molecules.^32^ Each system was solvated in a pre-equilibrated cubic box with a 12 Å buffer of TIP3P water molecules and was neutralized by addition of chloride counterions (Cl^-^). Subsequently, a two-stage geometry optimization approach was performed. First, a short minimization of water molecules, with positional restraints on solute by a harmonic potential with a force constant of 500 kcal mol^-1^ Å^-2^ was done. The second stage was an unrestrained minimization of all the atoms in the simulation cell. Then, the systems were heated using six steps of 50 ps, incrementing the temperature 50 K each step (0-300 K) under constant-volume, periodic-boundary conditions, and the particle-mesh Ewald approach to introduce long-range electrostatic effects.^33^ A 10 Å cut-off was applied to Lennard-Jones and electrostatic interactions. Bonds involving hydrogen were constrained with the SHAKE algorithm.^34^ Harmonic restraints of 10 kcal mol^-1^ were applied to the solute, and the Langevin equilibration scheme is used to control and equalize the temperature.^35^ The time step was kept at 2 fs during the heating stages. Each system was then equilibrated for 2 ns with a 2 fs timestep at a constant pressure of 1 atm to relax the density of the system. After the systems were equilibrated in the NPT ensemble, 50 replicas of 200 ns MD simulations for each protonation state of His57 (HID, HIE, HIP) were performed under the NVT ensemble and periodic-boundary conditions.

### Constant-pH Molecular Dynamics Simulations

A total of 30 replicas of 200 ns of spontaneous binding constant-pH MD simulations (CpH-MD) were run at pH = 7.0 and 8.0 considering residues His40, His57, and His91 as titratable.^13,14^ CpH-MD simulations were carried out using AMBER ff10 force-field in explicit TIP3P water. A number of 50 steps was assigned for the period of Monte Carlo steps and a salt concentration of 0.1 is assumed for the Generalized-Born calculation. A total of 100 relaxation steps were performed after a protonation state change.

## RESULTS

To evaluate the impact of His57 protonation state in the binding of benzamidine, we performed spontaneous binding molecular dynamics (MD) simulations placing four benzamidine molecules at least 30 Å away from the trypsin S1 pocket. In line with previous studies, we selected the structure corresponding to the trypsin-benzamidine complex (PDB ID 3PTB, see Methods) as starting point for the MD simulations. This implies that the trypsin S1 pocket is already predisposed for binding and, as shown by Buch et al., spontaneous binding can readily occur in the nanosecond time scale.^7^ From this starting point, benzamidine molecules are allowed to freely diffuse from the solvent and explore the trypsin surface in the predefined simulation time. Spontaneous binding MD simulations were performed using two different strategies. First, a total fifty replicas of 200 ns MD simulations were run for the three possible protonation states of His57: (1) delta nitrogen protonated (HID57); (2) epsilon nitrogen protonated (HIE57); and (3) positively charged histidine (HIP57). Second, we carried out a total of thirty replicas of 200 ns of constant-pH MD simulations at pH = 7.0 and 8.0 considering all histidine residues as titratable. From these simulations, the percentage of binding events and ligand pathways of benzamidine in trypsin were extracted. To evaluate the binding of benzamidine along the MD simulations, we monitored the distance between the carbon atom of the amidine group of benzamidine and the carbon of the carboxylate group of Asp189 side chain. We considered that productive binding is attained when this distance is below 5.4 Å, which is the minimum distance to retain the salt bridge interaction between the inhibitor and Asp189 in the S1 pocket.

### *The effect of His57 protonation states in benzamidine* binding

Our MD simulations showed that spontaneous binding of benzamidine to the S1 pocket can occur in all protonation states of His57 (see Figure 2). The preferred binding mode obtained from MD simulations in the three protonation states is equivalent to the one observed in the PDB 3PTB (trypsin-benzamidine complex), which is also the principal binding pose predicted in previous computational studies.^7,8,18^ However, a different number of binding events was observed for the three protonation states of His57. In particular, benzamidine binds the S1 pocket in 48% of HID57 simulations (24 out of 50 replicas of 200 ns), 44% of HIE57 (22 out of 50 replicas), and 10% of HIP57 (5 out of 50 replicas), as shown in Figure 2. Therefore, binding frequently occurs when His57 is found in its neutral form (either delta or epsilon nitrogen protonated) and becomes a less frequent event in its positively charged state (HIP57). This different distribution of binding events indicates that benzamidine binding is modulated by the protonation state of His57 which is located more than 10 Å away from Asp189. Considering only the replicas that captured productive binding events, the average binding times are: 88 ns, 80 ns, and 55 ns for HID57, HIE57, and HIP57 respectively. These results point out that binding can readily occur in all cases irrespective of the lower probability of binding events observed for HIP57. Therefore, the open question is how the different protonation states of His57 alter the probability of binding of benzamidine.

**Figure 2.**
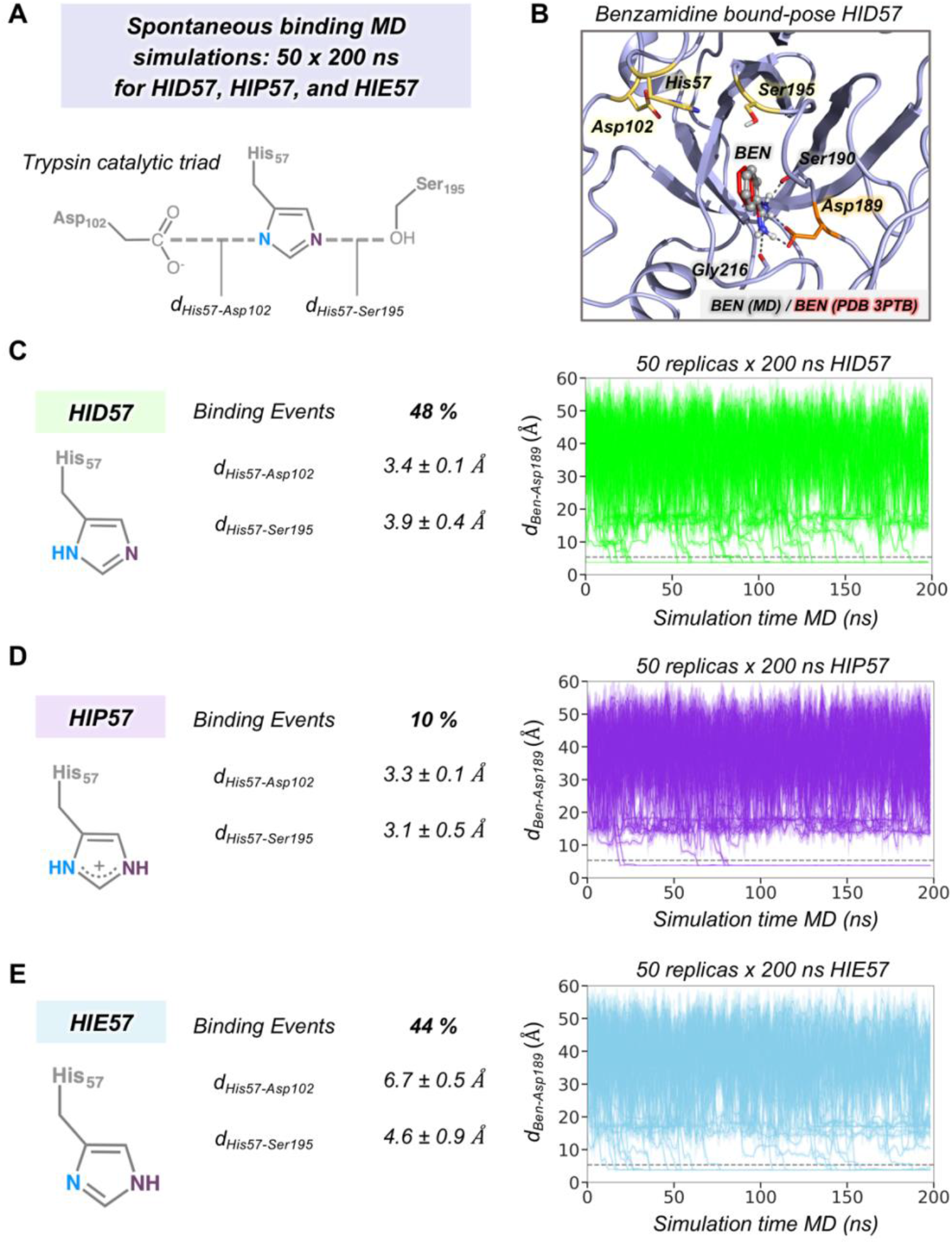
The effect of His57 protonation in benzamidine binding. (A) Representation of the trypsin catalytic triad formed by Asp102, His57, and Ser 195. The distance between Asp102 and His57 residues is calculated between the carbon of the carboxylate group of Asp102 and the delta nitrogen of His57. The distance between His57 and Ser195 is calculated between the epsilon nitrogen of His57 and the oxygen of the hydroxyl group of Ser195. (B) Representative conformation of the trypsin-benzamidine bound complex predicted from spontaneous binding HID57 MD simulations. The binding pose of benzamidine obtained from molecular dynamics (MD) simulations is shown in grey while the X-ray orientation (PDB 3PTB) is depicted in red. Catalytic residues and Asp189 are coloured in yellow and in orange, respectively. (C) Analysis of 50 replicas of 200 ns of HID57 MD simulations with the delta nitrogen of His57 protonated. (D) Analysis of 50 replicas of 200 ns of HIP57 MD simulations with His57 positively charged. (E) Analysis of 200 ns of HIE57 MD simulations with the epsilon nitrogen of His57 protonated. The percentage of binding events and the average distances (in Å) between catalytic residues is provided for each protonation state. Plot of the distance between the carbon atom of the amidine group of benzamidine and the carbon of the carboxylate group of Asp189 side chain along the 50 replicas of 200 ns MD simulations for each protonation state. The grey horizontal dashed line indicates when productive benzamidine binding takes place (distance below 5.4 Å).

In terms of global conformational dynamics, trypsin showed similar flexibility in the three protonation states of His57. Root-mean square fluctuations (RMSF) indicated that the flexibility of the S1 pocket is not significantly altered (see Figure S1). The main differences were observed when analyzing the stability of the catalytic triad formed by Asp102, His57, and Ser195 (see Figure 2). When His57 is positively charged (HIP57), the catalytic triad remains significantly stable, being 3.2 ± 0.5 Å and 3.3 ± 0.1 Å the Ser195-His57 and Asp102-His57 distances respectively. Additional flexibility is gained in the HID57 state with distances of 3.9 ± 0.4 Å (Ser195-His57) and 3.4 ± 0.1 Å (Asp102-His57). The intrinsic dynamism of the Ser195-His57 interaction in HID57 can be key to accommodate trypsin substrates triggering the formation of the Michaelis complex. Finally, the HIE57 protonation state is the least probable in trypsin because as shown in the MD simulations, when protonated in epsilon, His57 destabilizes the catalytic triad (see Figure 2). For this reason, all subsequent analyses will be focused on HID57 and HIP57 states.

### Characterization of Benzamidine Binding Pathways

To gain insight into the molecular basis of the effect of His57 protonation changes in the binding process, we explored the binding pathways that benzamidine followed to get into trypsin S1 pocket. First, we collectively represented all spontaneous binding MD simulations corresponding to each protonation state using two coordinates (see Figure 3): (1) the binding distance (*dBen-Asp189*, x axis) between the carbon atom of the amidine group of benzamidine and the carbon of the carboxylate group of Asp189; and (2) the distance between the carbon atom of the amidine group of benzamidine and the epsilon nitrogen of His57, for either HID57 or HIP57 (*d*_*Ben-His57*_, y axis). We selected the epsilon nitrogen because it is directly interacting with the hydroxyl group of catalytic Ser195. In both HID57 and HIP57, the most populated state of this 3D free-energy landscape (FEL) is the benzamidine-bound state (i.e. *d*_*Ben-Asp189*_ below 5 Å and *d*_*Ben-His57*_ around 10 Å, see **B** in Figure 3). Benzamidine also accumulates in both cases in a region of the FEL defined by a range of *d*_*Ben-Asp189*_ [15,20] Å and *d*_*ben-His57*_ [10,15] Å, which corresponds to trypsin hydrophobic S3 pocket composed by Trp210 and surrounding residues. Interestingly, the FEL displays significant differences in the binding patterns of HID57 and HIP57 when benzamidine approaches the S1 pocket (*d*_*Ben-Asp189*_ within [5,15] Å). For HID57, the FEL shows a metastable state where benzamidine directly interacts with His57 (*d*_*ben-His57*_ below 5 Å while *d*_*Ben-Asp189*_ is still found between 10 and 15 Å, see **I**_**1**_ and **I**_**2**_ states in Figure 3A). This state is not visited in the FEL of HIP57 indicating that the interaction between His57 and benzamidine is not established in these MD simulations. These results are not surprising considering that both benzamidine and His57 are positively charged in HIP57 simulations resulting in repulsive interactions that prevent the interaction. From these simulations, we calculated the free-energy difference between the unbound conformation and the transition state that leads to productive binding (see Figure S2). The free-energy difference is 2.3 kcal/mol and 3.8 kcal/mol for HID57 and HIP57 respectively, pointing out that the binding of benzamidine is globally slowed down when His57 is positively charged.

**Figure 3.**
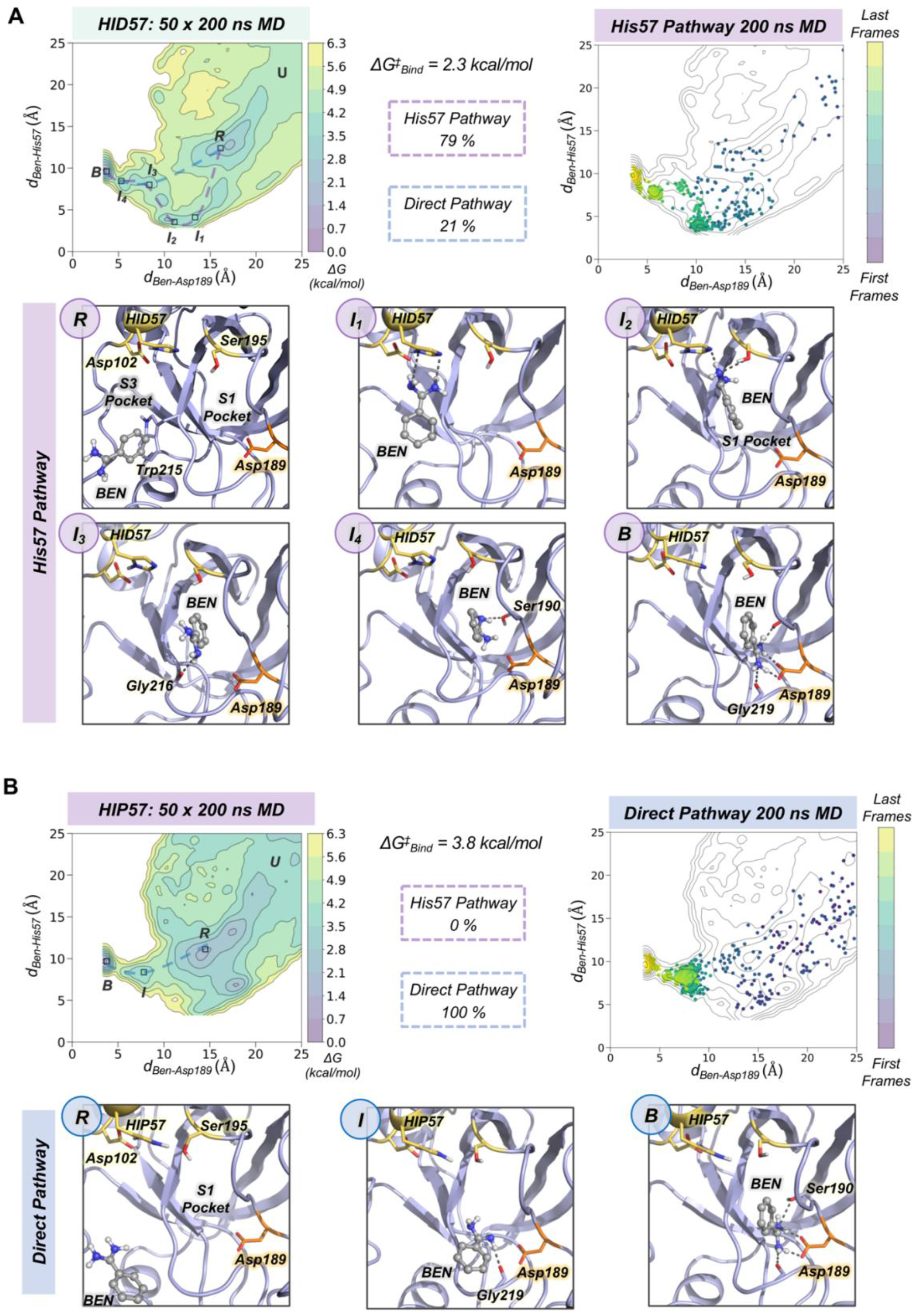
Characterization of benzamidine binding pathways. Free energy landscape (FEL) reconstructed from 50 replicas of 200 ns of spontaneous binding MD simulations of HID57 (A) and HIP57 (B) using the binding distance (*d*_*Ben-Asp189*_, x axis) between the carbon atom of the amidine group of benzamidine and the carbon of the carboxylate group of Asp189 and the distance between the carbon atom of the amidine group of benzamidine and the epsilon nitrogen of His57 (*d*_*Ben-His57*_, y axis). The most relevant states of the FEL are highlighted in black boxes: *U* (unbound), *R* (recognition), *B* (bound) and *I* (intermediate) states. The free-energy difference between the unbound conformation and the transition state that leads to productive binding is given in kcal/mol. The purple dashed line indicates the trajectory of the His57 pathway while the blue dashed line indicates the trajectory of the direct pathway. The percentage of binding events (considering only productive binding simulations) that follow each pathway is provided. Projection of a representative spontaneous binding 200 ns MD trajectory on the FEL of each HID57 and HIP57. The time evolution of the ligand binding pathway is represented in a colour scale ranging from purple for the first frames to yellow for the last frames of the MD trajectory. Molecular representation of the most relevant states of the FEL corresponding to the His57 and direct pathways. Catalytic residues are shown in yellow, benzamidine in grey, and Asp189 in orange.

To characterize the molecular basis of the ligand binding processes, we independently analyzed the MD trajectories projecting them into the corresponding FEL (see Figures 3 and S3). The analysis of independent HID57 MD trajectories showed that benzamidine binding occurred mainly through two different pathways. In the major binding pathway (observed in 79% of productive binding simulations), benzamidine first enters the S3 pocket being recognized by Trp210. Then, the inhibitor rolls along this pocket to establish a hydrogen bond interaction with His57 while keeping the phenyl moiety in the S3 pocket. Then, the aromatic ring of benzamidine repositions from the S3 to the S1 pocket maintaining the hydrogen bonding with catalytic His57 and Ser195. Finally, through a series of successive steps, benzamidine turns around to establish a salt bridge interaction with the carboxylate group of Asp189, attaining the benzamidine-bound pose. We identify this predominant pathway as “His57 pathway” since establishing a hydrogen bond interaction with His57 is a requisite to access the S1 pocket. A similar binding pathway was described by Buch and co-workers using 495 100 ns MD simulations.^7^ A second, less frequent, pathway was observed in 21 % of productive binding simulations. In this case, benzamidine directly evolves from the solvent to the S1 pocket without establishing a hydrogen bond interaction with HID57 (see Figure 3). The direct access to the S1 pocket from the solvent can take place from different sites but the common feature is that benzamidine directly enters the pocket establishing a hydrogen bond with Ser190 and then repositions to interact with Asp189 through a salt-bridge interaction. In this particular pathway, the binding of benzamidine is practically a pure diffusion from the solvent to the S1 pocket, for this reason we identify it as the “direct pathway”.

In HIP57 MD simulations, benzamidine binding only occurred through the direct pathway (5 out of 50 replicas), as shown in Figure 3B. The repulsion between the positive charges of both protonated His57 (HIP) and benzamidine prevents their approximation and interaction. Thus, the His57 pathway is not observed in HIP57 simulations, making the number of binding events significantly less frequent than in HID57. Despite benzamidine can be recognized also in the S3 pocket by Trp210, when it approaches the positively charged HIP57 a hydrogen bond with this catalytic residue cannot be established and benzamidine returns back to the solvent (see Figure S4). Therefore, when His57 is positively charged, binding will preferentially occur through direct diffusion from the solvent. These results demonstrate that the protonation state of His57 determines which ligand binding pathways can be populated and, thus, the path that benzamidine preferentially takes to access the S1 pocket of trypsin.

### Spontaneous Benzamidine Binding with Constant-pH Molecular Dynamics Simulations

Using fixed protonation states for His57 can offer a limited picture of benzamidine binding, considering that this residue is responsible for the most relevant pKa shifts during binding and catalysis in trypsin.^20^ To account for the pH effects in the benzamidine binding process, we performed thirty replicas of 200 ns of spontaneous binding constant-pH MD simulations at pH = 7.0 and 8.0 (see Figures 4 and S5). In these simulations, we allowed the three histidines of trypsin to change their protonation state. Only His57 is found along the binding pathway and can play a key role in the benzamidine recognition process. At pH = 7.0, the populations of the different protonation states obtained from CpH-MD simulations were 74%, 26%, and 0% for HIP, HID, and HIE, respectively. From these CpH-MD simulations at pH = 7.0, we obtained a FEL of the binding process that resembles the one captured for HIP57 protonation state (see Figures 3B and 4A). However, the intermediate states corresponding to the interaction between benzamidine and His57 became moderately populated. This increases the number of binding events compared to HIP57 simulations up to 23% (7 out of 30 replicas). In this case, benzamidine accessed the S1 pocket through both direct and His57 pathways. Interestingly, CpH-MD simulations showed that the His57 pathways is activated only when His57 attains the HID protonation state (see Figure 4A). At pH = 8.0, we observed equilibrium populations of protonation states for His57 of 57% for HID, 42% for HIP, while HIE is scarcely populated. The FEL obtained from CpH-MD simulations at pH = 8.0 clearly highlights the stabilization of the interaction between His57 and benzamidine (see **I**_**1**_ and **I**_**2**_ states in Figure 4B). Again, binding of benzamidine proceeded through both the direct and His57 pathways. The number of binding events retrieved from CpH-MD simulations at both pH = 7.0 and 8.0 is found between the values observed in HIP57 and HID57 MD simulations offering a clearer picture of the His57 protonation state ensemble. Therefore, properly accounting for the equilibrium of protonation states of His57 may be key to retrieve accurate kinetics for the binding of benzamidine to trypsin.

**Figure 4.**
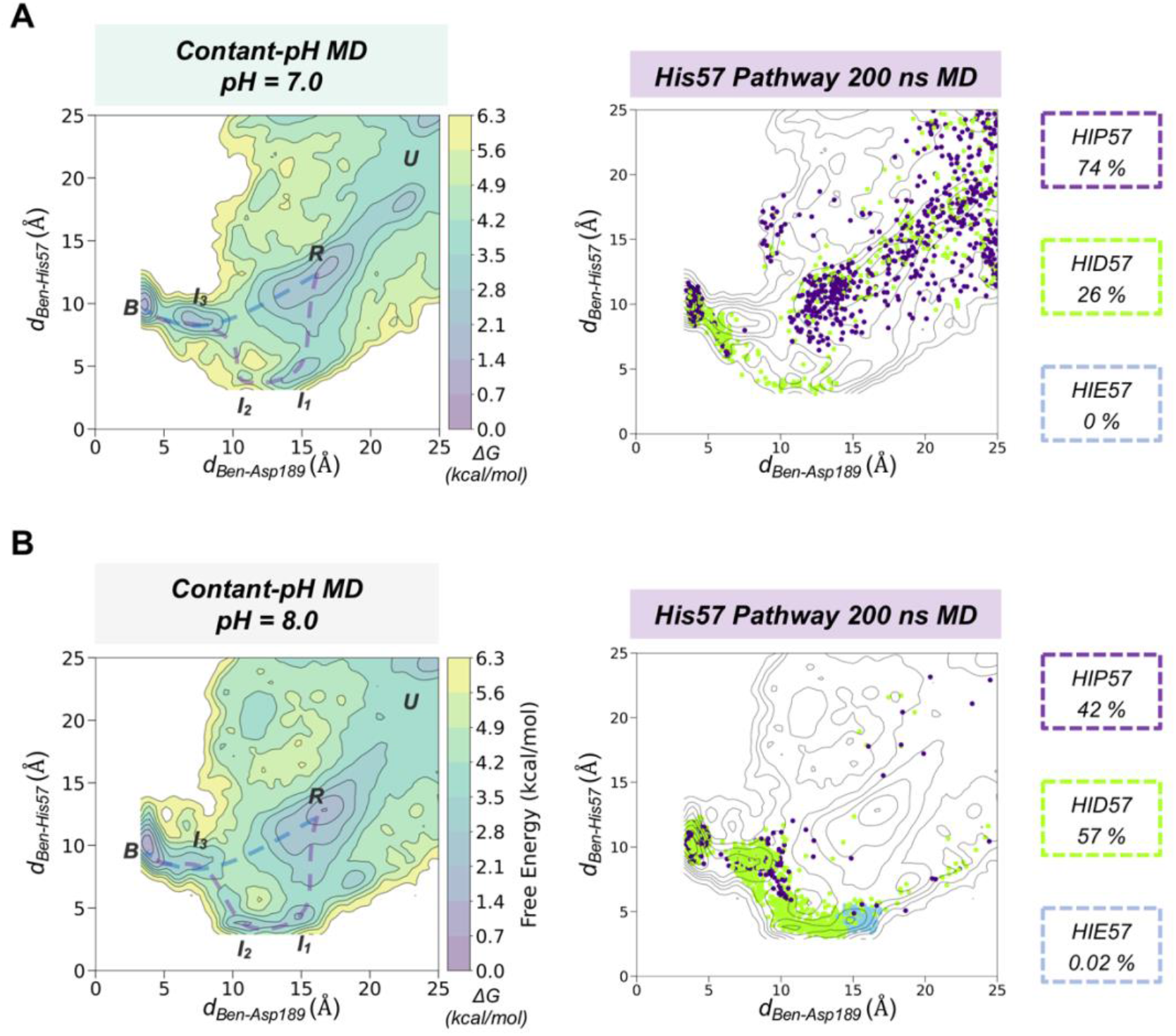
Spontaneous Benzamidine Binding with Constant-pH Molecular Dynamics Simulations. Free energy landscape (FEL) reconstructed from 30 replicas of 200 ns of spontaneous binding constant-pH MD simulations at pH = 7.0 (A) and pH = 8.0 (B) using the binding distance (*d*_*Ben-Asp189*_, x axis) between the carbon atom of the amidine group of benzamidine and the carbon of the carboxylate group of Asp189 and the distance between the carbon atom of the amidine group of benzamidine and the epsilon nitrogen of His57 (*d*_*Ben-His57*_, y axis). The most relevant states of the FEL are highlighted in black boxes: *U* (unbound), *R* (recognition), *B* (bound) and *I* (intermediate) states. Projection of a representative 200 ns trajectory of the His57 pathway showing the protonation state of each frame in different colour: HID in green, HIP in purple, and HIE in blue. The equilibrium populations of each protonation state retrieved from the 30 replicas of 200 ns is provided for pH = 7.0 and 8.0.

## DISCUSSION

Spontaneous binding MD simulations showed that the protonation state of His57, which is located more than 10 Å away from the gorge of the S1 pocket, plays a key role in determining the binding pathway of benzamidine to trypsin. Binding is more favorable when His57 attains a neutral HID protonation state while is less probable in the positively charged HIP protonation. Benzamidine can access the S1 pocket through two main pathways: the His57 pathway and the direct pathway from the solvent. We observed that His57 is found in the way of benzamidine to the S1 pocket through the most probable binding pathway in the HID protonation state, establishing a hydrogen bond that is key to drive benzamidine toward the binding site. Therefore, subsequent kinetic analysis of the binding process will provide different outcomes depending on how the protonation states are defined at the beginning of the simulation. Constant-pH MD simulations naturally account for the protonation state ensemble of His57 offering a more accurate description of the spontaneous binding of benzamidine at a fixed pH.

Based on these results, we suggest to always consider the impact of protonation changes of residues that are found along the ligand binding pathway (even distal residues) when performing spontaneous binding MD simulations. In particular, in systems with slow binding processes that follow complex pathways through different metastable intermediate and transition states. It is important to remark that in our study we have excluded protonation changes of other residues (e.g. Asp, Glu, …) which may further alter the binding pathways obtained. These observations are not limited to the study of drug-binding into their biological receptors. In enzyme engineering, substrate access tunnels are commonly engineered through point mutations to evolve the enzyme toward a new function or broad its substrate scope. Thus, properly accounting for coupled protonation state changes between the residues conforming the access tunnel and the substrate will be important to evaluate the substrate binding pathways in enzymes. By unravelling the details of ligand binding and unbinding it is possible to gain insight into the detailed molecular mechanisms of relevant biochemical processes and then, harness this information to rationally improve the potency of drugs and/or evolve an enzyme toward novel functions.

## ACKNOWLEDGEMENTS

We thank Carla Calvó-Tusell and Dr. Miguel A. Maria-Solano for fruitful discussions and Daniel Masó for technical assistance. We thank Dr. Yinglong Miao for helpful discussions on spontaneous binding simulations of the trypsin-benzamidine complex. This work was supported by the Spanish MICINN projects PID2019-111300GA-I00 (M.G.B.), RTI2018-101032-J100 (F.F). We thank Spanish MICINN for Ramón y Cajal fellowships RYC2020-028628-I (M.G.B.) and RYC2020-029552-I (F.F.). We thank the Generalitat de Catalunya for the emerging group CompBioLab (2017 SGR-1707), the consolidated group DiMoCat (2017 SGR-39), for predoctoral FI fellowship 2022 FI_B 00615 (H.G.) and Beatriu de Pinós H2020 MSCA-Cofund 2018-BP-00204 project (to M.G.B.).

## Notes

### Competing Interest Statement

The authors have declared no competing interest.

