## Supplementary Information for "Changes in protonation states of in-pathway residues can alter ligand binding pathways obtained from spontaneous binding molecular dynamics simulations"

*A. Root Mean Square Fluctuations (RMSF) Trypsin: HID 57, HIP57, HIE57*

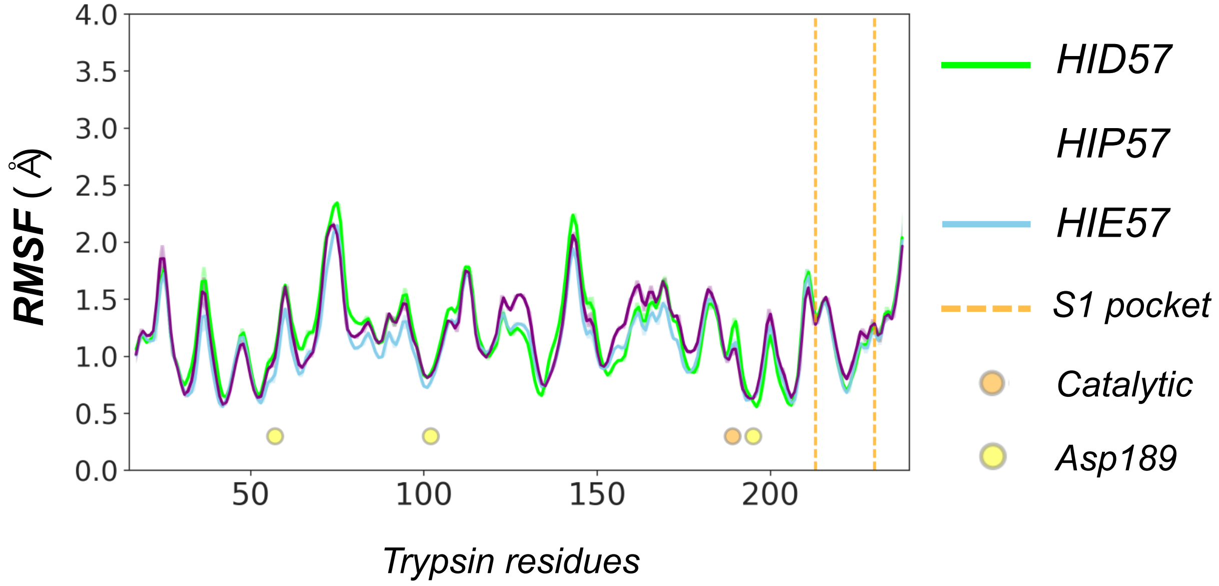


**Supplementary Figure 1.** Plot of the Root-Mean-Square-Fluctuation (RMSF, in Å) for trypsin HID57 (green), HIDP57 (purple), and HIE57 (blue) obtained from fifty replicas of 200 ns MD simulations. The catalytic residues are highlighted in yellow and Asp189 in orange. Vertical orange dashed lines indicate the position of the S1 pocket loop. In terms of global flexibility, all systems present a similar flexibility.

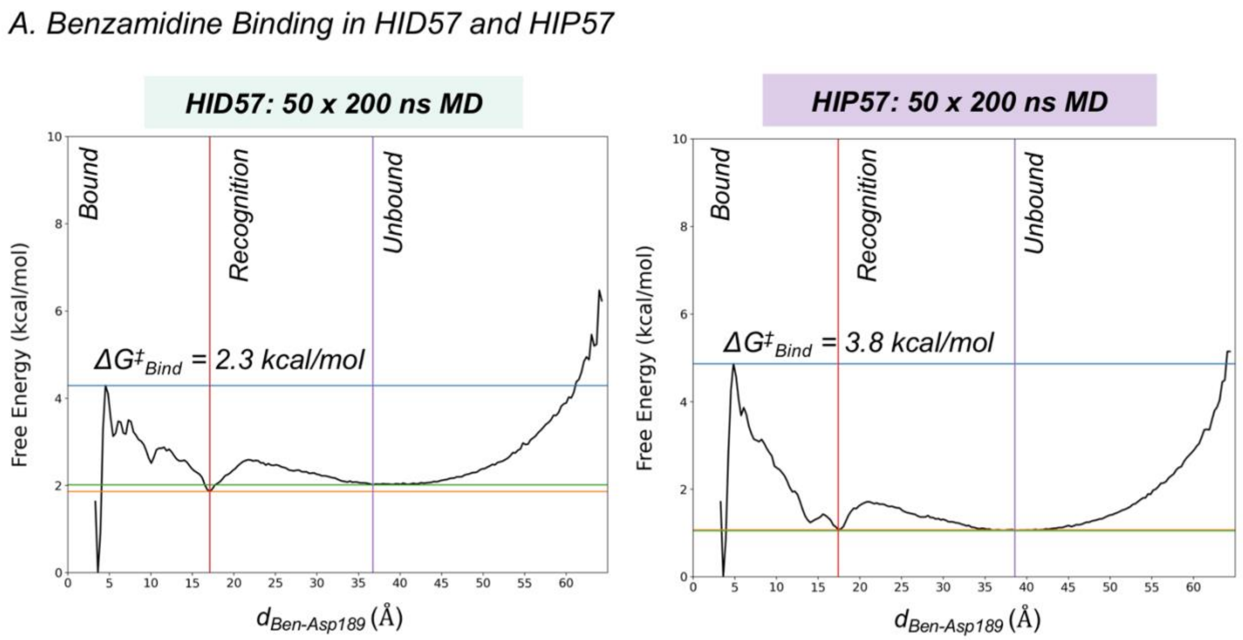


**Supplementary Figure 2.** (A) Free energy profile reconstructed from 50 replicas of 200 ns of spontaneous binding MD simulations of HID57 and HIP57 using the binding distance ($d_{Ben\text{-}Asp189}$, x axis) between the carbon atom of the amidine group of benzamidine and the carbon of the carboxylate group of Asp189 and the distance between the carbon atom of the amidine group of benzamidine and the epsilon nitrogen of His57 ($d_{Ben\text{-}His57}$, y axis). The free-energy difference ($\Delta G^{\ddagger}_{Bind}$) between the unbound conformation and the transition state that leads to productive binding is provided for each case. The free-energy difference is 2.3 kcal/mol and 3.8 kcal/mol for HID57 and HIP57, respectively pointing out that the binding of benzamidine is globally slowed down when His57 is positively charged.

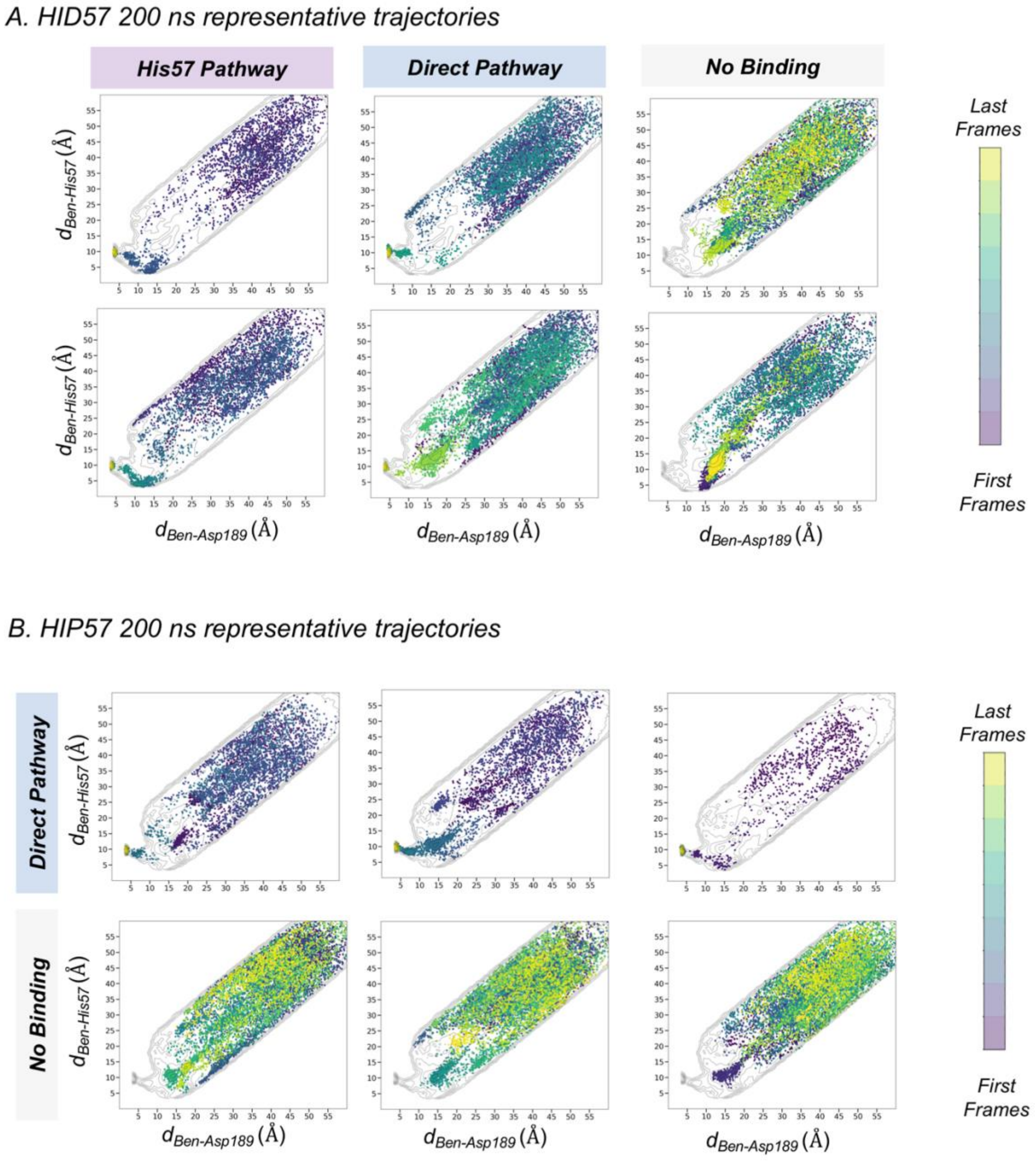


**Supplementary Figure 3.** Projection of representative spontaneous binding 200 ns MD trajectory on the FEL (see Figure 3 main text) of each HID57 and HIP57. Free energy landscape (FEL) reconstructed from 50 replicas of 200 ns of spontaneous binding MD simulations of HID57 (A) and HIP57 (B) using the binding distance ($d_{Ben\text{-}Asp189}$, x axis) between the carbon atom of the amidine group of benzamidine and the carbon of the carboxylate group of Asp189 and the distance between the carbon atom of the amidine group of benzamidine and the epsilon nitrogen of His57 ($d_{Ben\text{-}His57}$, y axis). Selected examples of the His57 pathway, direct pathway, and replicas where binding is not observed are provide for each protonation state. The time evolution of the ligand binding pathway is represented in a colour scale ranging from purple for the first frames to yellow for the last frames of the MD trajectory.

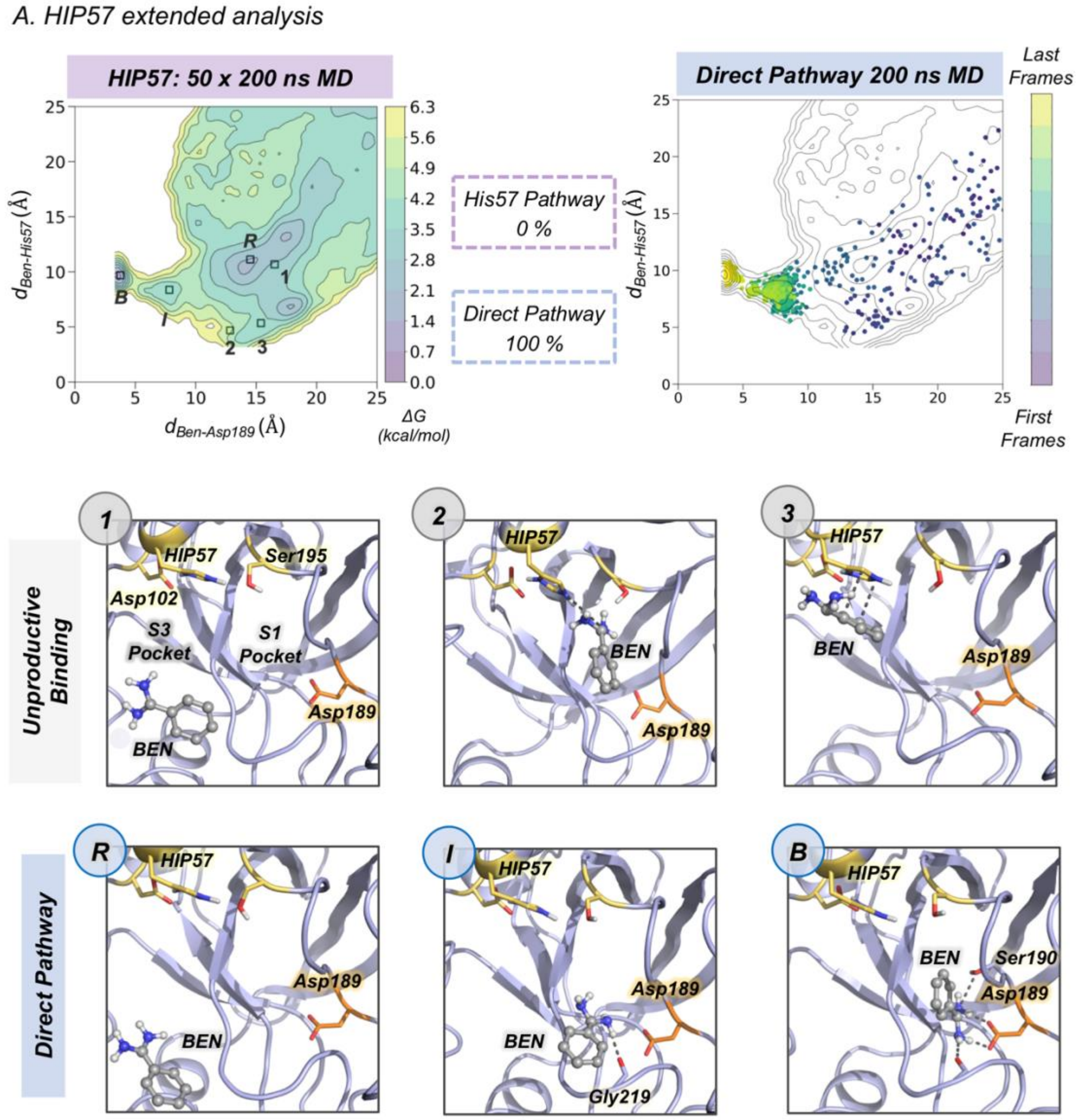


**Supplementary Figure 4.** (A) Free energy landscape (FEL) reconstructed from 50 replicas of 200 ns of spontaneous binding MD simulations of HIP57 using the binding distance ($d_{Ben\text{-}Asp189}$, x axis) between the carbon atom of the amidine group of benzamidine and the carbon of the carboxylate group of Asp189 and the distance between the carbon atom of the amidine group of benzamidine and the epsilon nitrogen of His57 ($d_{Ben\text{-}His57}$, y axis). The most relevant states of the FEL are highlighted in black boxes: *U* (unbound), *R* (recognition), *B* (bound) and *I* (intermediate) states while unproductive binding pathways are highlighted in humber. The percentage of binding events (considering only productive binding simulations) that follow each pathway is provided. Projection of a representative spontaneous binding 200 ns MD trajectory on the FEL of HIP57. The time evolution of the ligand binding pathway is represented in a colour scale ranging from purple for the first frames to yellow for the last frames of the MD trajectory. Molecular representation of the most relevant states of the FEL corresponding to the direct pathways. Catalytic residues are shown in yellow, benzamidine in grey, and Asp189 in orange. When benzamidine approaches His57 repulsive interactions between positively charges His57 and benzamidine bring the inhibitor back to the solvent.

A. Constant-pH MD simulations at pH = 7.0: representative trajectories

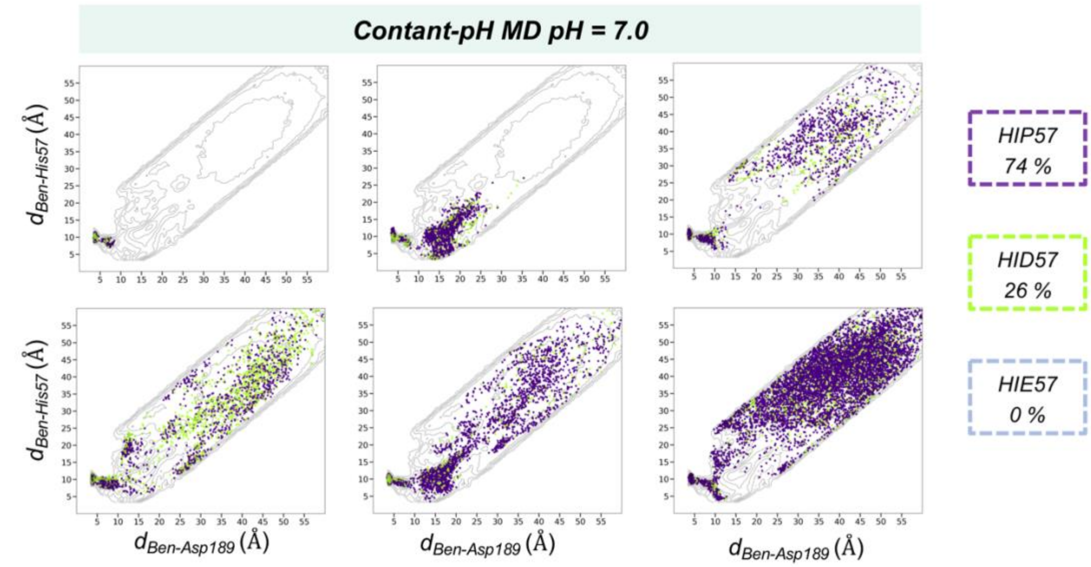


B. Constant-pH MD simulations at pH = 8.0: representative trajectories

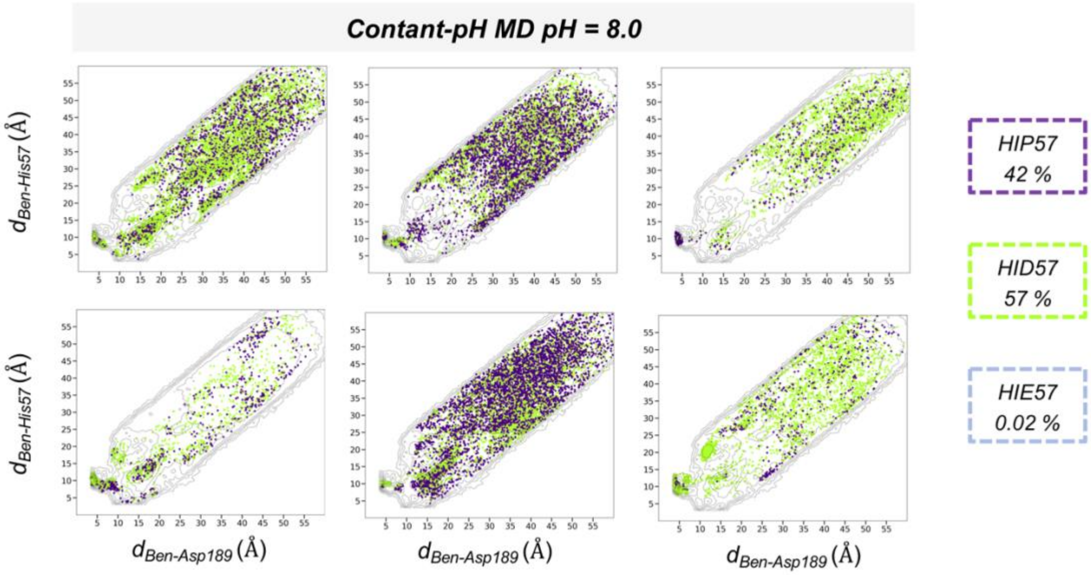


**Supplementary Figure 5.** Free energy landscape (FEL) reconstructed from 30 replicas of 200 ns of spontaneous binding constant-pH MD simulations at pH = 7.0 (A) and pH = 8.0 (B) using the binding distance ($d_{Ben\text{-}Asp189}$, x axis) between the carbon atom of the amidine group of benzamidine and the carbon of the carboxylate group of Asp189 and the distance between the carbon atom of the amidine group of benzamidine and the epsilon nitrogen of His57 ($d_{Ben\text{-}His57}$, y axis). The most relevant states of the FEL are highlighted in black boxes: *U* (unbound), *R* (recognition), *B* (bound) and *I* (intermediate) states. Projection of a representative 200 ns trajectory of the His57 pathway showing the protonation state of each frame in different colour: HID in green, HIP in purple, and HIE in blue. The equilibrium populations of each protonation state retrieved from the 30 replicas of 200 ns is provided for pH = 7.0 and 8.0.
